# Approximation of bone mineral density and subcutaneous adiposity using T1-weighted images of the human head

**DOI:** 10.1101/2024.05.22.595163

**Authors:** Polona Kalc, Felix Hoffstaedter, Eileen Luders, Christian Gaser, Robert Dahnke

## Abstract

Bones and brain are intricately connected and scientific interest in their interaction is growing. This has become particularly evident in the framework of clinical applications for various medical conditions, such as obesity and osteoporosis. The adverse effects of obesity on brain health have long been recognised, but few brain imaging studies provide sophisticated body composition measures. Here we propose to extract the following bone- and adiposity-related measures from T1-weighted MR images of the head: an approximation of skull bone mineral density (BMD), skull bone thickness, and two approximations of subcutaneous fat (i.e., the intensity and thickness of soft non-brain head tissue). The measures pertaining to skull BMD, skull bone thickness, and intensi-ty-based adiposity proxy proved to be reliable (*r*=.93/.83/.74, *p*<.001) and valid, with high correlations to DXA-derived head BMD values (rho=.70, *p*<.001) and MRI-derived abdominal subcutaneous adipose volume (rho=.62, *p*<.001). Thickness-based adiposity proxy had only a low retest reliability (*r*=.58, *p*<.001).The outcomes of this study constitute an important step towards extracting relevant non-brain features from available brain scans.

## Introduction

The brain is closely connected to other tissues and organs (Zhou et al. 2023), including the skeletal system. A large body of literature suggests a bidirectional communication between bones and the brain (Obri et al. 2018; Rousseaud et al. 2016). Moreover, bone-derived metabolites have been implicated in mood, cognition, and glucose/energy homeostasis in murine models (Guntur & Rosen 2012; Khrimian et al. 2017; Lee et al. 2007; Nakamura et al. 2021). However, the neuroimaging community largely dismisses the information from the closest bone structure surrounding the brain.

Bone tissue undergoes constant remodelling which is supported by osteoclasts (i.e., bone resorbing cells) and osteoblasts (i.e., new bone-forming cells) to regulate mineral homeostasis and ensure skeletal integrity (Clarke 2008; Feng and McDonald 2011). Bone metabolism is affected by endocrine signalling and innervation via the sympathetic nervous system, with greater sympathetic activity leading to decreasing BMD (Ducy and Kousteni 2015; Takeda et al. 2002). Furthermore, bone tissue actively signals to the brain and other organs and is implicated in brain metabolism by secretion of bone-derived peptides (e.g., osteocalcin, osteopontin, sclerostin, lipocalin-2, etc.), as well as by production of bone marrow-derived immune cells (Chen et al. 2021; Yu et al. 2020).

The relation between BMD and fat mass is complex (Lee et al. 2020; Rinonapoli et al. 2021). For example, people with anorexia nervosa tend to have lower BMD than healthy and obese individuals (Fazeli & Kliban-ski 2014; Maïmoun et al. 2020). However, increased adiposity levels or other metabolic disorders can can have a detrimental effect on bone health, as the mesenchymal stromal cells of the bone marrow in obese people preferentially convert into adipocytes (i.e. fat cells) rather than osteoblasts (i.e. new bone-forming cells), potentially resulting in reduced bone mass (Ambrosi et al. 2017; Veldhuis-Vlug & Rosen 2018). Bone health also seems to play a role in neurodegenerative as well as neurodevelopmental disorders (Kelly et al., 2020), such as Parkinson’s disease (Sleeman, Che, and Counsell 2016; Tassorelli et al. 2017), multiple sclerosis (Bisson et al. 2019), and autism spectrum disorder (Neumeyer et al. 2015). Moreover, a low BMD has been associated with an increased risk of Alzheimer’s Disease (Kostev et al. 2018; Lui et al. 2003; Tan et al. 2005; Zhang et al. 2022; Zhou et al. 2011; Xiao et al. 2023).

Given the consistent reports on the connection of bone measures (e.g., BMD, bone mineral content) and brain ageing in the UK Biobank database (e.g., Miller et al. 2016; Smith et al. 2019; 2020) as well as the long-known adverse effects of obesity on cognitive and brain health (Beydoun et al. 2008; Farruggia & Small 2019), the availability of such measures in open-source brain imaging databases, such as IXI, ADNI, AIBL, or OASIS is inadequate. Here we present the extraction and validation of skull BMD, skull bone thickness, and non-brain soft head tissue thickness and intensity estimation from the head tissue classes, normally discarded when processing T1-weighted brain MR images.

## Methods

### Data

This research has been conducted using data from the UK Biobank (UKB application #41655), a biomedical database and research resource that contains genetic, lifestyle and health information from half a million UKB participants (https://www.uk-biobank.ac.uk/). The UKB has ethical approval from the North West Multi-Centre Research Ethics Committee (MREC) and is in possession of informed consents from the study cohort participants.

T1-weighted brain MR images of a subsample of healthy UKB subjects (N = 2000, M_Age_ = 64.17±6.34 years, age range: 46–89 years, 50% women) were used to extract and validate our skull BMD estimate against BMD measure of the head and total body BMD obtained by dual energy X-ray absorptiometry (DXA) scanning. Our head tissue intensity and thickness measures (i.e., subcutaneous fat approximation) was evaluated with respect to body mass index, percent body fat, abdominal subcutaneous adipose tissue volume (ASAT), visceral adipose tissue volume (VAT), and waist circumference.

Furthermore, we used a sample of 316 T1-weighted MR scans from OASIS-3 dataset, acquired at two time points within a time interval of less than 3 months (M_Age_ = 67.66 ± 8.22 years, age range: 42–85, 56% women) to determine the retest reliability of the measures for the same as well a (slightly) different scanners and protocols. To evaluate the original and our refined bone segmentation, a subsample of individuals with both MR and CT scans was used (N_CT_ = 114, M_Age_ = 66.71 ± 8.38 years, age range: 42–85, 51% women).

The quality of the scans was monitored internally through the UKB workflow (Alfaro-Almagro et al., 2018), and was assessed by the authors in OASIS-3.

### Data acquisition

#### Bone mineral density data acquisition

BMD measures of the head and total body were obtained by dual energy X-ray absorptiometry (DXA) scanning with iDXA instrument (GE-Lunar, Madison, WI). A detailed overview of the acquisition procedure is available in https://biobank.ndph.ox.ac.uk/showcase/ukb/docs/DXA_explan_doc.pdf.

#### Quantification of the subcutaneous adipose tissue

Subcutaneous adipose tissue in the abdomen was acquired by abdominal MRI imaging on Siemens MAGNETOM Aera 1.5 T MRI scanner (Siemens Healthineers, Erlangen, Germany) with a 6-minute dual-echo Dixon Vibe protocol, resulting in a water and fat separated volumetric data set (Linge et al. 2018) that was processed by AMRA Profiler Research (AMRA Medical AB, Linköping, Sweden) to obtain body composition measures (Borga et al. 2015).

#### Brain MRI data acquisition

All the information regarding brain MR data acquisition is available in the UK Biobank Imaging documentation (https://biobank.ctsu.ox.ac.uk/crystal/crystal/docs/brain_mri.pdf). In short, T1-weighted MR images were acquired on Siemens Skyra 3.0 T scanners with a 32-channel RF receive head coil. The MPRAGE sequence was used with 1-mm isotropic resolution, inversion/repetition time = 880/2000 ms, acquisition time: 5 minutes, FOV: 208x256x256 matrix, in-plane acceleration factor = 2.

The MRI data from the OASIS-3 dataset used in this study were acquired on different 1.5 T and 3.0 T scanners with 20-channel head coil (Siemens Sonata, Siemens TIM Trio and BioGraph mMR PET-MR 3T) (LaMontagne et al., 2019). The MPRAGE sequence was used with (i) 1 mm isotropic repetition/echo time = 1900/3.93 ms, FA = 15° FOV phase = 87.50%, (ii) 1 mm isotropic inversion/repetition/echo time = 1000/2400/3.16 ms, FA = 8°, FOV phase = 100%, R = 2 and (iii) 1.20x1.05x1.05 mm resolution, inversion/repetition/echo time = 900/2300/2.95 ms, FA = 9°, FOV phase = 93.75 mm, R = 2.

#### CT scan data acquisition

CT scans were obtained on the Siemens Biograph 40 PET/CT scanner. A three-second X-ray topogram was acquired in the lateral plane and a spiral CT scan was performed for attenuation correction at the low dose CT (LaMontagne et al. 2019).

### Data preprocessing

Raw T1-weighted MR brain images were processed using SPM12 (Wellcome Center for Human Neuroimaging, https://www.fil.ion.ucl.ac.uk/spm/) running under Matlab 2021a (Mathworks Inc., Natick, MA, USA), which produced the following segments: grey matter, white matter, cerebrospinal fluid, skull, soft head tissue, and background. Of note, instead of the default setting (3 mm), we set SPM’s ‘samp’ parameter to 5 mm to ensure that non-brain tissues were properly classified.

The CT data was preprocessed with CTseg utilising the unified segmentation of SPM12 (Brudfors et al., 2020; https://github.com/WCHN/CTseg) and co-registered to the individual MRI space.

### Development of the measures

#### Bone measures

The SPM processing often erroneously assigned the high intensity bone marrow to the head tissue class instead of the skull segment. Mis-classified skull segments were therefore corrected by morphological operations to avoid missing the intensities from the diploë (i.e., cancellous (spongy) bone between the inner and outer layer of the cortical bone of the skull) and to correct the underestimated bone thickness (Figure 2B).

**Figure 1.**
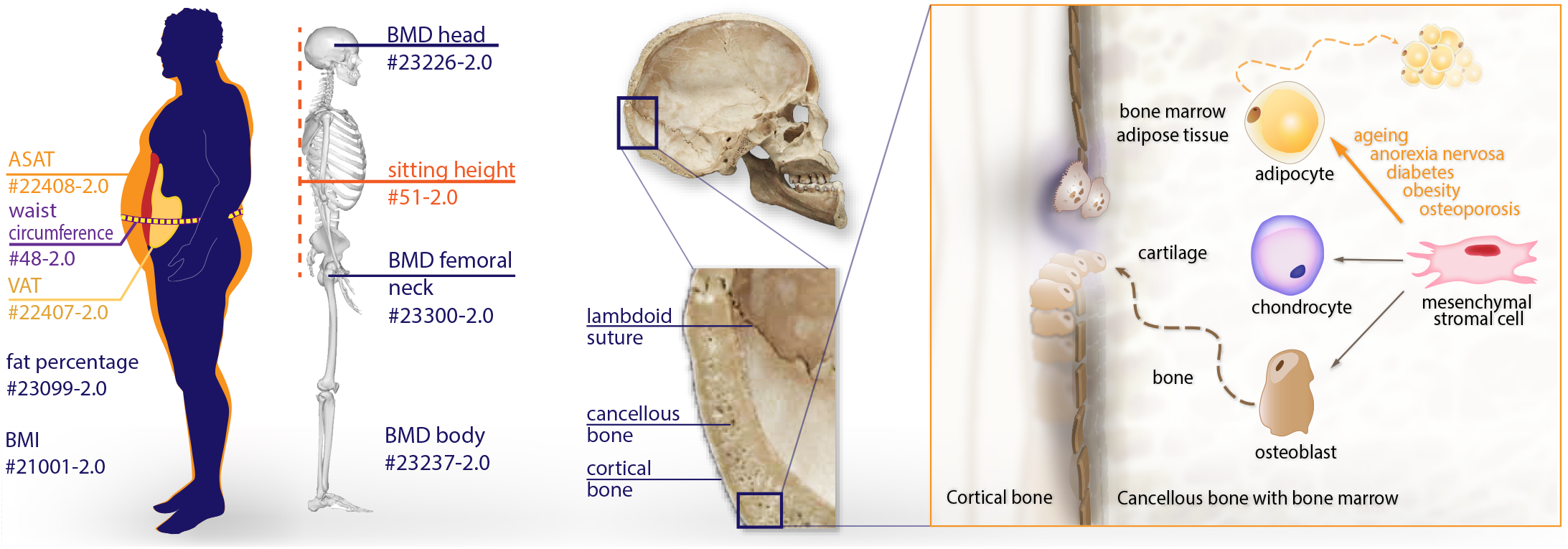
Demonstrates the UKB anthropometric measures used in the validation process as well as the cranial anatomy and microenvironment of a bone. Bone marrow is located within the spongy/cancellous bone (diploë). Mesenchymal stromal cells in the bone marrow have the potential to develop into osteoblasts (bone-forming cells), chondrocytes (cartilage-forming cells), or adipocytes (bone marrow fat cells). Obesity, ageing, diabetes, anorexia nervosa, starvation, and osteoporosis lead their differentiation into adipocytes and, hence, a lower BMD (Tencerova & Kassem 2016).

**Figure 2.**
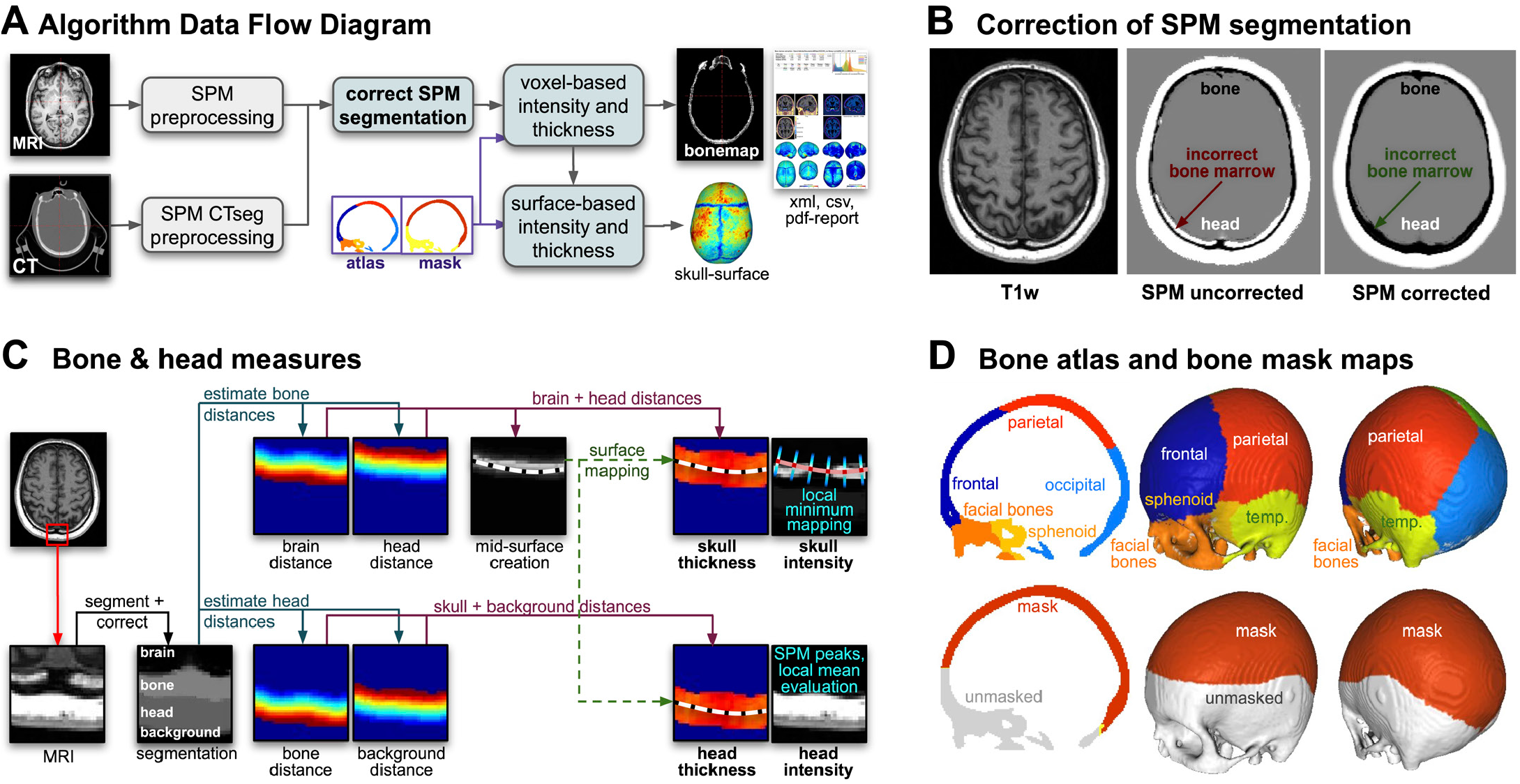
(A) The figure shows the workflow of the algorithm with necessary corrections (B), extraction of measures (C), and template maps (D). MRI/CT images undergo preprocessing by SPM/SPM-CTseg into CSF, GM, WM, skull, head, and background. Morphological operations are used to correct segmentation errors (B), before estimating various distance maps to describe the skull and head tissue thickness maps (C). The calvarial atlas and mask, shown in (D), are mapped to the individual space to extract regional and global values. If surface processing is required, the distance maps are used to create a percentage position map, which allows the creation of the mid-skull-surface. This surface is then used to map the local thickness, intensity and atlas region to the surface to extract a regional or masked global thickness and intensity value.

**Figure 3.**
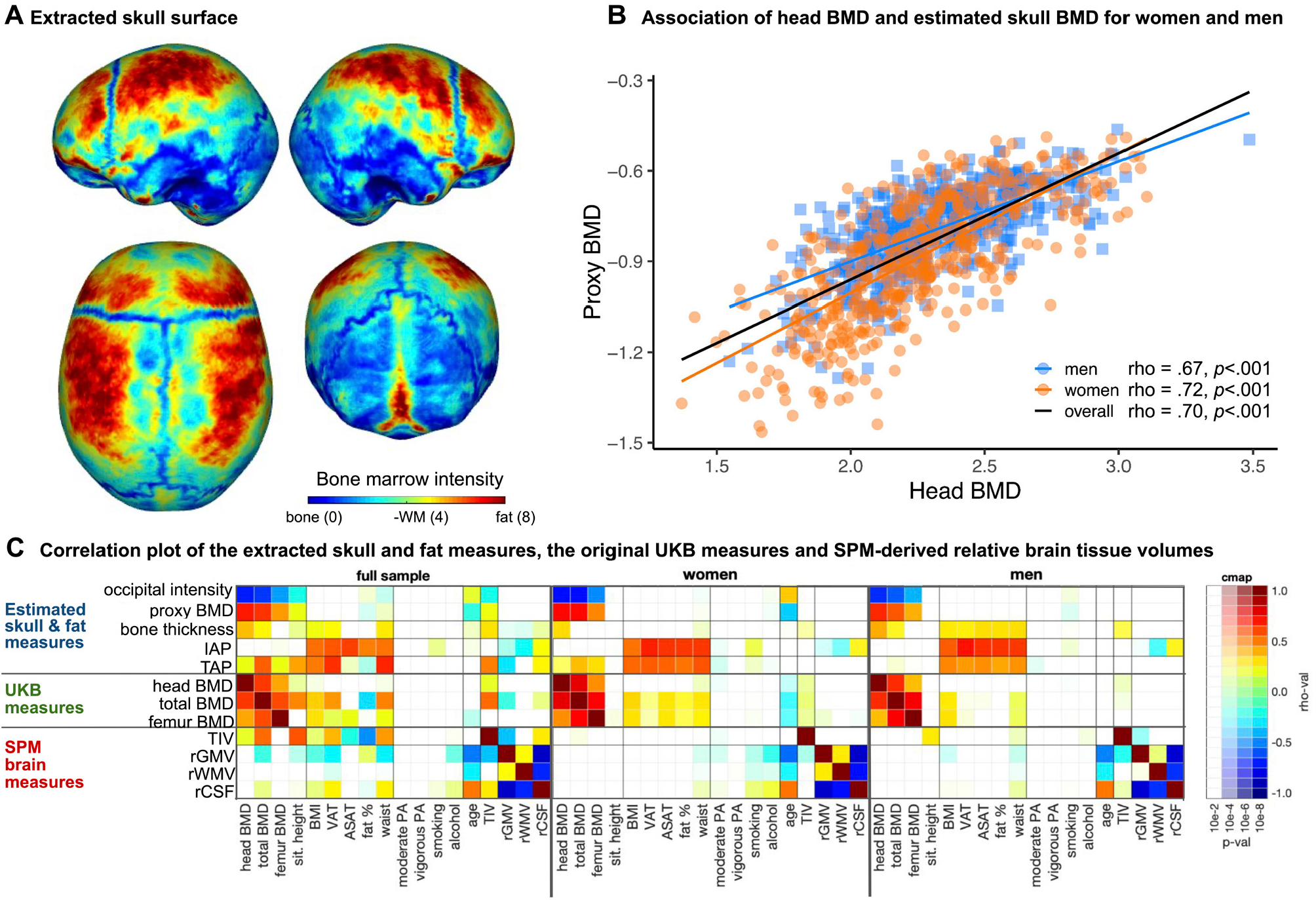
(A) The extracted skull surface of a subject from the UK Biobank with visible sutures and high intensity bone marrow. (B) Scatterplot of the UKB head BMD and our skull BMD estimate for men and women. Higher head BMD is related to lower (occipital) bone intensity in the MRI and therefore results in a higher proxy BMD value. (C) Spearman correlation coefficients with Holm correction for multiple comparisons between the estimated skull BMD proxy, bone thickness, as well as intensity- and thickness-based adiposity proxies (IAP & TAP, respectively), and other UKB and brain measures. The extracted measures are associated with the original BMD as well as body composition measures.

##### Intensity-based measures

Bone measures were derived from the corrected SPM skull segment, by quantifying the mean intensity across the entire segment labelled as skull (or within a specific skull region), and normalised by the SPM-derived WM intensity.

##### Thickness-based measures

Bone thickness was defined as the sum of the shortest distance from each skull voxel to the CSF and to the soft non-brain head tissue. In addition to voxel-based processing, a midline surface was generated to map bone values along the surface-normal. This allowed the cortical bone intensity, quantified as the local minimum, to be separated from the bone marrow intensity, defined as the distance-weighted average of the mapped points (Figure 2C).

#### Head tissue measures

##### Intensity-based measures

To approximate the adipose tissue of the head, we estimated as a weighted average of the SPM-derived Gaussian peaks by their proportions in the tissue class labelled as head. Head tissue intensity was described as a percentage composite of the various soft non-brain head tissues.

##### Thickness-based measures

Local head thickness was calculated as the shortest distance from each soft head tissue voxel to the skull and to the background. The sum of both distances then yielded the estimate of the vox-el-wise head tissue thickness. A separation into different head tissues (i.e., muscle, skin, and fat) was omitted because of varying amounts of (chemical shift) artefacts and inhomogeneities across the image. Of note, the lower portions of all brain scans as well as voxels located more than 30 mm from the skull were excluded from the head tissue thickness estimation to avoid side effects due to defacing and/or varying scanning protocols and procedures.

#### Boney Atlas

The processing results yielded clearly identifiable coronal, sagittal and lambdoid sutures, visibly separating the calvarial bones. We therefore created a template atlas of the skull regions by averaging affine and intensity-normalised T1-weighted images as well as CT data of OASIS-3, and manually labelling the skull segment tissue probability map using Slicer3D (Fedorov et al. 2012; https://slicer.readthedocs.io/en/5.0/index.html). The atlas was mapped into individual space using the linear transformation from the SPM segmentation. Additionally, we created a bone mask to avoid critical regions that are often affected by defacing or are in parts very thin, such as temporal and sphenoid bones (Figure 2D). The identified bone regions were used in extracting regional bone and adiposity measures.

### Validation of the measures

The segmentation results were visually examined against ground truth CT data, juxtaposing MR- and CT-derived measures from the OASIS-3 dataset (see Figure S1 in the Supplement”“).

Global and regional measures were tested to obtain the best estimates of the BMD in an exploratory manner on a subsample of 1000 subjects (M_Age_ = 64.02 ± 6.40 years, age range: 46–80 years, 50% women). A random forest regression with all the extracted bone measures as independent variables predicting the UKB head’s BMD measure and a permutation test were run in R (version 4.3.3; R Core Team 2022) with packages *party* (Hothorn et al., 2006) and *flexplot* (Fife, 2022). The final BMD approximation measure was chosen based on the estimated predictor importance values from a permutation test (Fife & D’Onofrio, 2023).

The selected measures were validated by calculating the Spearman correlation coefficient (*psych* package; Revelle, 2024) to relevant measures available in the UKB in a subsample of 1000 subjects not included in the exploratory analysis (M_Age_ = 63.81 ± 6.27 years, age range: 46–79 years, 50% women). We calculated the associations with head BMD (UKB #23226-2.0) and total body BMD (UKB #23237-2.0), as well as body fat percentage (UKB #23099-2.0) and abdominal subcutaneous adipose tissue (UKB #22408-2.0) volume, for the estimates of skull BMD and head tissue thickness, respectively (see Figure 1““).

We additionally inspected the correlation to other anthropometric and lifestyle variables that had previously been connected to BMD or adiposity (Cusano, 2015; Molenaar et al. 2009; Shojaa et al. 2020; Ward & Klesges, 2001; Williams et al., 2005). We investigated the association with BMI (UKB #21001-2.0), sitting height (UKB #51-2.0), visceral adipose tissue volume (VAT; UKB #22407-2.0), waist circumference (UKB #48-2.0), and BMD of the left femoral neck (UKB #23300-2.0). Furthermore, we included the variables connected to physical activity, namely duration of moderate (UKB #894-2.0) and vigorous physical activity (UKB #914-2.0), as well as pack years of smoking as proportion of life span exposed to smoking (UKB #20162-2.0) and frequency of alcohol intake (UKB #1558-2.0).

As an additional validation step, we performed a volume-based morphometry (VBM) analysis in SPM12 with BMD of the head and our newly extracted proxy BMD as predictors of the extracted warped and smoothed (FWHM = 8 mm) intensity-normalised skull segments on the whole sample of 2000 UKB subjects.

### Evaluation of reliability

We calculated the retest reliability of the estimated measures using T1-weighted images from the OASIS-3 dataset acquired at two time points within an interval of less than 3 months. One subject was excluded from the analysis due to a failed SPM-segmentation, resulting in a total sample of 157 subjects (age range: 42–85, M_Age_ = 67.64 ± 8.24 years, 55% women). The reliability of the measures was estimated under the same scanner/protocol and mixed scanner/protocol conditions.

## Results

### Validation of the measures on the UK Biobank dataset

The results of the exploratory step showed that the most relevant measure for the BMD approximation is the mean intensity extracted from the occipital bone (Table S1 in the Supplement<u>““</u>). The results were confirmed by the VBM analysis of the skull segments, where we could see the highest association of the head BMD and the occipital region of the skull (Figure 4““). We assumed the same pattern for the head tissue thickness and intensity estimates based on visual inspection of the images. All further analyses therefore included the occipital measures to represent the skull BMD, skull bone thickness, intensity-based, and thickness-based adiposity proxies.

**Figure 4.**
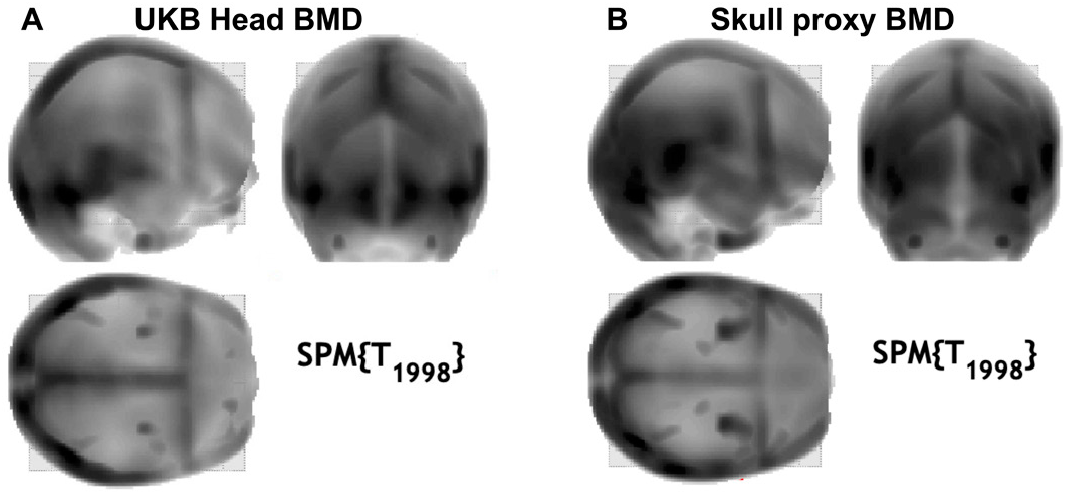
‘Glass’ head maps from the VBM analysis in SPM with (A) head BMD from the UKB as a predictor of the intensity-normalised skull segments, and (B) proxy BMD measure as a predictor of the intensity-normalised skull segments.

The validation results against other anthropometric and lifestyle measures are shown in Figure 3C (and Figure S2 in the Supplement““). The extracted measures showed valid associations to other UKB parameters. The Spearman’s correlation coefficients between the extracted proxy BMD measure and UKB-derived head BMD were .70, .72, and .67 for a total sample, women, and men, respectively (see Figure S3““ in the Supplement for the bootstrapped correlation coefficient). The correlation with the total body BMD was moderate for the total sample and the subsample of men (rho = .64 and .57, respectively, *p* <.0001) and higher for women (rho = .72, *p* <.0001). The association to other anthropometric measures showed a weak negative correlation with total fat percent (rho = –.14, *p* <.01), and a low positive correlation with waist circumference (rho = .15, *p* <.01). A weak negative association between the proxy BMD measure and the exposure to smoking was observed in a subsample of men (rho = –.12, *p* <.01). Bone thickness (in the occipital part) was, on the other hand, weakly associated with body composition measures in men (ASAT: rho = .31, *p* <.0001; waist circumference: rho = .31, *p* <.0001, percent body fat: .30, *p* <.0001).

Our intensity-based adiposity approximation was moderately to highly positively correlated with other body composition measures. The Spearman’s correlation coefficients with BMI were .52, .52, and .57 for a total sample, women, and men, respectively. The association is higher for a volume of visceral adipose tissue (VAT), namely .57, .68, and .70 for a total sample, women, and men, respectively, and abdominal subcutaneous adipose tissue volume (ASAT; rho = .62; rho = .64; rho = .65, for a total sample, women, and men, respectively). Similar patterns were observed in our thickness-based adiposity approximation. The measure was moderately positively associated with other body composition measures. The Spearman’s correlation coefficients with BMI were .50, .54, and .48 for a total sample, women, and men, respectively. The association with VAT volume was higher, namely .65, .56, and .45 for a total sample, women, and men, respectively. The thickness-based adiposity proxy had a similar correlation with VAT as BMI in a total sample (rho = .65, *p* < .0001; rho = .64, *p* < .0001 for VAT association with head thickness and BMI, respectively), but lower in subsamples of men and women (see Figure S2 in the Supplement”“).

It could be observed that the measures were biassed due to anthro-pometric differences between the biological sexes. For subsequent use in statistical analyses, the sex, age and TIV of the participants must be taken into account.

### The reliability of the measures (OASIS-3)

The retest reliability based on the data obtained with the same protocol on two different time-points (n = 63, M_Age_ = 69.27 ± 7.32 years, 48-85, 54% women) was high for the skull BMD (r = .93; *p* < .001) and bone thickness estimation (r = .83, *p* < .001), comparatively to the regional GM volume (r = .98, *p* < 0.001). Our thickness-based adiposity proxy showed lower retest reliability (r = .58, *p* < .001). Nevertheless, the intensity-based adiposity proxy had a sufficient reliability of .74 (*p* < .001). It should be noted that the retest reliability of the measures derived from images acquired with different protocols is generally lower than for identical protocols. The results from the whole selected sample (n = 157) and varying protocols between the two time-points (n = 94, M_Age_ = 66.91 ± 8.81 years, 42–85, 50% women) are available in the Supplementary Table S2”“.

## Discussion

In the present study, we used tissue classes that are typically discarded when processing brain images to estimate skull BMD, skull bone thickness, as well as intensity- and thickness-based adiposity approximations. Such measures are normally not considered in neuroimaging research. However, burgeoning research shows the interconnectedness between brain, bones, and adipose tissue (Caron et al., 2018; Rousseaud et al., 2016). Thus, the aim of this new line of research is to provide the neuroimaging community with proxy measures that can be extracted from standard T1-weighted brain scans that are available in open-source databases or that have been acquired by researchers following standard protocols of brain imaging.

### Reliability and validity of measures

All extracted measures showed a good retest reliability on the OASIS-3 dataset in both same- and mixed-protocol test cases, with the only exception of thickness-based adiposity proxy. Its low retest reliability might be due to bias introduced during the head segment correction, and further development is currently underway to improve this specific measurement.

The validity of the proxy BMD measure was confirmed by moderate to high associations with DXA-derived BMD measures of the head and total body. The proxy BMD measure also showed a link to age in women (but not men). This is expected as women experience significant changes in bone mass with ageing due to decreases in oestrogen and increases of follicle-stimulating hormone (Almeida, 2012; Iqbal et al. 2012). Men, on the other hand, are less susceptible to age-related hormonal fluctuations, and their bone mass tends to remain stable until later in life (Porter & Varacallo, 2022). However, certain medications and poor lifestyle choices (e.g., physical inactivity or smoking) can also increase the risk of developing osteoporosis in men (Porter & Varacallo, 2022). In fact, our proxy BMD measure was partly linked to smoking exposure in men.

The validity of skull bone thickness was confirmed by association with certain aspects of body composition measures. Higher BMI and abdominal adiposity were weakly related to higher skull bone thickness in the occipital region. These results could be linked to the body weight-related mechanical stress to the bone (Cherukuri et al. 2021). However, it is typically the lean body mass that has a positive effect on bone health, rather than fat mass (Ho-Pham et al., 2014, Nguyen et al., 2020). Nevertheless, the connection between skull bone thickness and adiposity might involve the effects of adipokines (i.e., cytokines produced by the adipocytes) on bone mass (Mangion et al. 2023) and further studies are needed to elucidate this finding.

The associations of thickness- and intensity-based adiposity proxy measures with UKB anthropometric measures are moderate to high; BMI had the lowest correlation and waist circumference and VAT the highest. This lower connection to BMI is not surprising, as BMI is inherently a proxy measure of body composition (Gutin, 2018). Among other drawbacks, BMI fails to account for changes in body composition when ageing, where lean muscle mass typically decreases while visceral adiposity increases, resulting in no change in weight (Bhurosy & Jeewon 2013). In an ageing sample like the UKB, such limited association to BMI is expected. Nevertheless, a stronger association between thickness-based adiposity proxy with VAT (in comparison to BMI) is crucial, as it shows the relevance of its extraction for the approximation of abdominal obesity. Moreover, both adiposity proxy measures were also negatively correlated with the relative GM volume and positively with CSF volume, which corroborates the previous findings on the negative influence of obesity on brain structure (Gómez-Apo et al., 2021).

### Potential application

Previous studies have shown that head fat tissue volume is linked to estimates of body composition in people with obesity (Wang et al. 2014). Nevertheless, to our knowledge, we conducted the first study that provides an estimate of BMD from a T1-weighted scan of the head. The outcomes of our study suggest that skull BDM can serve as a proxy measure of a person’s total BMD. In addition, the information on skull bone thickness might become relevant in studies of traumatic brain in-juries as the cranium plays a vital role in protecting the brain (Semple & Panagiotopoulou, 2023). Furthermore, the thickness-based adiposity proxy measures can be included as potential confounds in functional neuroimaging studies, such as electroencephalography, functional near infrared spectroscopy, and transcranial direct current stimulation (Gor-niak et al. 2022). Last but not least, the adiposity proxy measures may provide valuable information beyond the typically used BMI, which has several limitations (Burkhauser & Cawley 2008; Tomiyama et al. 2016), especially when used in cohorts of older individuals (Bhurosy & Jee-won 2013; Rothman 2008). This might be particularly relevant as many open-source brain imaging databases (e.g., ADNI, AIBL, OASIS) include a high percentage of adults who are past midlife.

### Limitations and future directions

The bone and adiposity estimations derived from T1-weighted MRI scans are not suitable for clinical use. This is especially relevant as structural MRI protocols can vary in fat-suppression intensity, which can affect the estimates in various ways. Characterising the nature of these variations and developing robust estimations for MRI protocols with fat suppression is warranted in future studies. Additionally, our proxy BMD measure is highly correlated with head BMD, but only moderately with the BMD of femoral neck, which is a better predictor of fracture risk (Kanis et al. 2004). With its high morbidity and mortality in older adults and its association to AD, fracture risk is a relevant aspect of healthy ageing (Friedman et al., 2010). Thus, future studies utilising machine learning might focus on establishing a better predictor of (femoral neck) BMD. Moreover, deep learning might aid in segmenting specific tissue classes, such as muscles, skin, and fat, which could result in an even better (skinfold-like) approximation of an individual’s status related to overall health as well as obesity (Langner et al. 2020; Leong et al. 2024). Altogether, the present study serves as a pilot exploring the potential value of non-brain tissue analyses in the field of neuroimaging. Further studies are necessary to improve and validate the measures.

## Data and code availability statement

The data used for this work were obtained from the UK Biobank Resource (Project Number 41655). Due to the nature of the data sharing agreement, we are not allowed to publish the data. The OASIS-3 dataset is openly available at https://www.oasisbrains.org. Prior to accessing the data, users are required to agree to the OASIS data use terms (DUT), which follow the creative commons attribution 4.0 licence. The code developed in this study is available on GitHub: https://github.com/robdahn/boney

## Authorship contribution statement

**Polona Kalc:** Conceptualization, Formal analysis, Validation, Writing – original draft, Writing – review and editing, Visualisation. **Felix Hoffstaedter:** Resources, Data curation, Writing – review & editing. **Eileen Luders:** Writing – review & editing. **Christian Gaser:** Supervision, Funding acquisition, Writing – review & editing. **Robert Dahnke:** Conceptualization, Software, Data curation, Validation, Visualisation, Writing – review & editing.

## Acknowledgements

This research has been conducted using data from UK Biobank under the application number 41655. We gratefully acknowledge the time and effort of the UK Biobank participants as well as the researchers and staff involved in data collection and management.

Data were provided in part by OASIS-3: Longitudinal Multimodal Neuroimaging: Principal Investigators: T. Benzinger, D. Marcus, J. Morris; NIH P30 AG066444, P50 AG00561, P30 NS09857781, P01 AG026276, P01 AG003991, R01 AG043434, UL1 TR000448, R01 EB009352. AV-45 doses were provided by Avid Radiopharmaceuticals, a wholly owned subsidiary of Eli Lilly.

## Supplementary material

**Figure S1.**
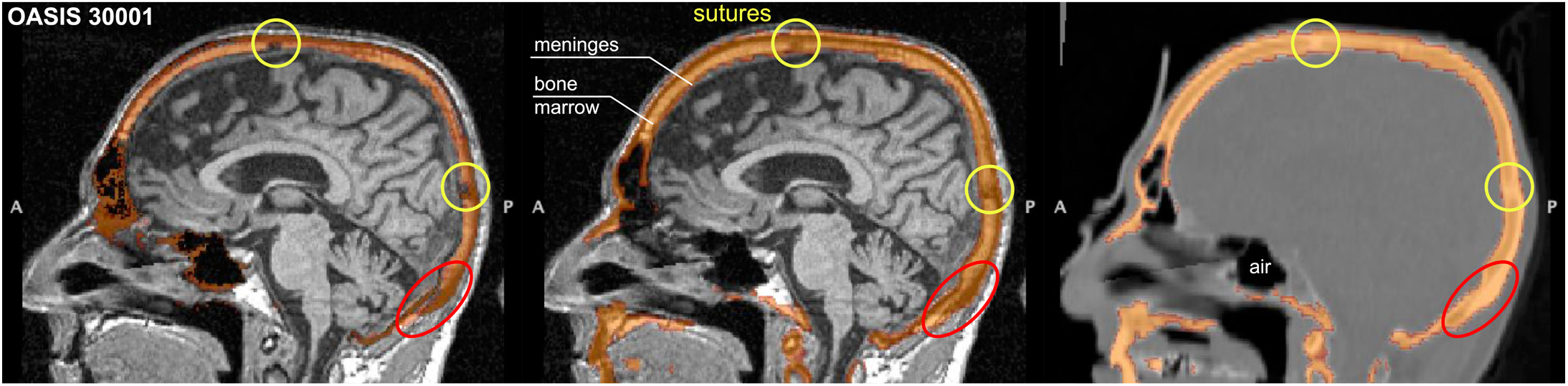
(A) Segmentation result of an OASIS-3 subject for MRI and CT. Overlap of the original SPM MRI segmentation (left) and the CTseg CT segmentation on the MRI (middle) and the co-registered CT images. The general overlap of the MRI and CT image is very good. The oversegmention of the skull on the top of the head could indicate light chemical shift artefacts. Details of the bone structure are visible in both, CT and MRI, where sutures (yellow circles) are darker in MRI but brighter in CT, and the bone marrow is brighter in MRI but darker in CT (also in case of fat suppression) (red).

**Table S1.** Results of the best predictor estimation in 5 folds.

| Fold | Best predictor 1<br>[variable importance] | Best predictor 2<br>[variable importance] | Best predictor 3<br>[variable importance] | Variance explained by the<br>model |
| --- | --- | --- | --- | --- |
| 1 | sROI_bonecortex01<br>0.053 | **main_sBMDH**<br>0.051 | sROI_bonethickness03<br>0.051 | 0.62 |
| 2 | **sROI_bonecortex03**<br>0.052 | vROI_bonethickness01<br>0.047 | sROI_bonecortex01<br>0.045 | 0.60 |
| 3 | **sROI_bonecortex03**<br>0.052 | **main_sBMDH**<br>0.051 | sROI_bonethickness03<br>0.043 | 0.60 |
| 4 | **sROI_bonecortex03**<br>0.061 | **main_sBMDH**<br>0.056 | sROI_bonecortex01<br>0.055 | 0.65 |
| 5 | **main_sBMDH**<br>0.054 | sROI_bonecortex04<br>0.051 | **sROI_bonecortex03**<br>0.051 | 0.63 |

Notes. sROI\_bonecortex01: surface-based global bone intensity value, sROI\_bonecortex03: surface-based occipital bone intensity value, sROI\_bonecortex04: surface-based right parietal bone intensity value; sROI\_bonethickness03: thickness of the occipital bone; main\_sBMDH is the opposite value of the sROI\_bonecortex03 (mean occipital bone intensity value); Variable importance: difference in RMSE of the model using predicted vs. permuted observations (Fife & D'Onofrio, 2023).

**Table S2.** Pearson’s correlation coefficient for OASIS-3 rescans within 3 months (p < 0.001). The images were mostly acquired on different Siemens scanners with slightly different protocols independent of the time point which introduce variations comparable to independent sites.

| Measures | Full sample<br>(N = 157) | Same protocol<br>(N = 63) | Mixed protocols<br>(N = 94) |
| --- | --- | --- | --- |
| Occipital bone intensity (sROI_bonecortex03) | 0.71 | **0.93** | 0.71 |
| Occipital bone thickness (sROI_bonethickness03) | 0.81 | **0.83** | 0.79 |
| SPM head tissue class intensity (tis_head) | 0.67 | **0.74** | 0.59 |
| Occipital head thickness (sROI_headthickness03) | 0.45 | **0.58** | 0.51 |
| Relative GM volume (rGMV) | 0.94 | **0.98** | 0.90 |

**Figure S2.**
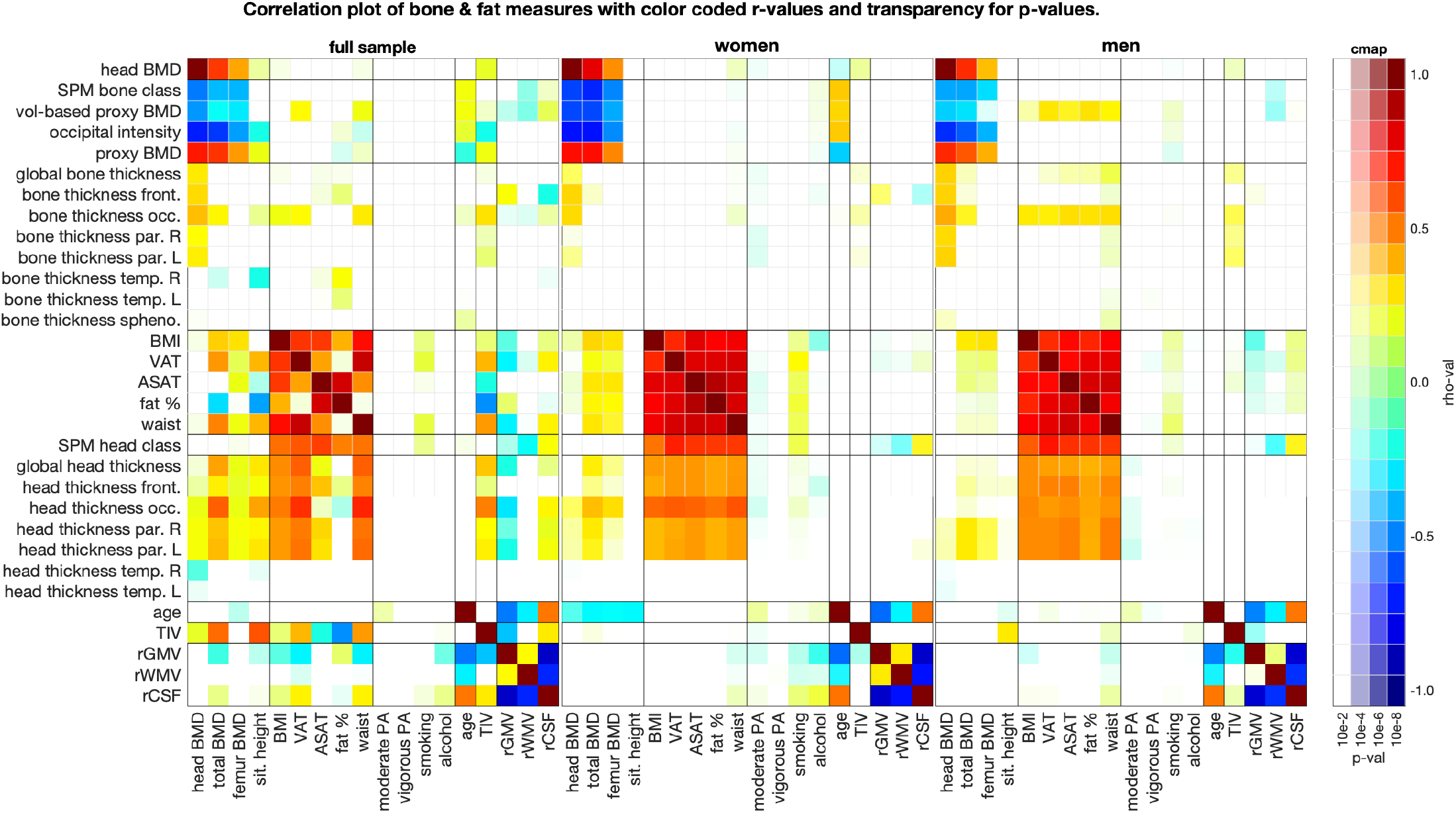
Spearman correlation coefficients with Holm correction for multiple comparisons between the estimated skull BMD proxy, head thickness, and other UKB and brain measures. SPM bone class represents a weighted average of the 3 Gaussian curves estimated by the SPM in the bone class. Vol-based proxy BMD is the volume-derived estimation of the bone intensity in the occipital bone. Proxy BMD is the opposite value of the surface-derived estimation of the bone intensity in the occipital bone.

**Figure S3.**
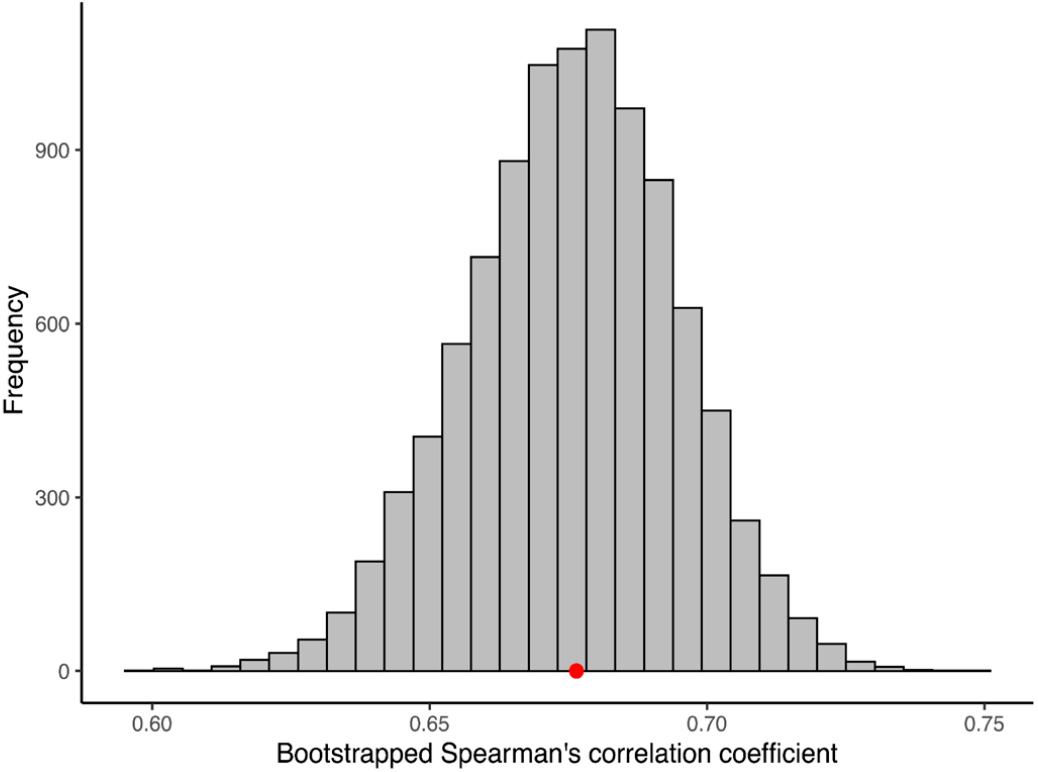
A distribution of bootstrapped Spearman’s correlation coefficients between the proxy BMD measure and DXA-derived head BMD. The red dot represents the Bootstrap estimated correlation coefficient (rho = 0.68, p<.001, 95% CI 0.64–0.71).

**Figure S4.**
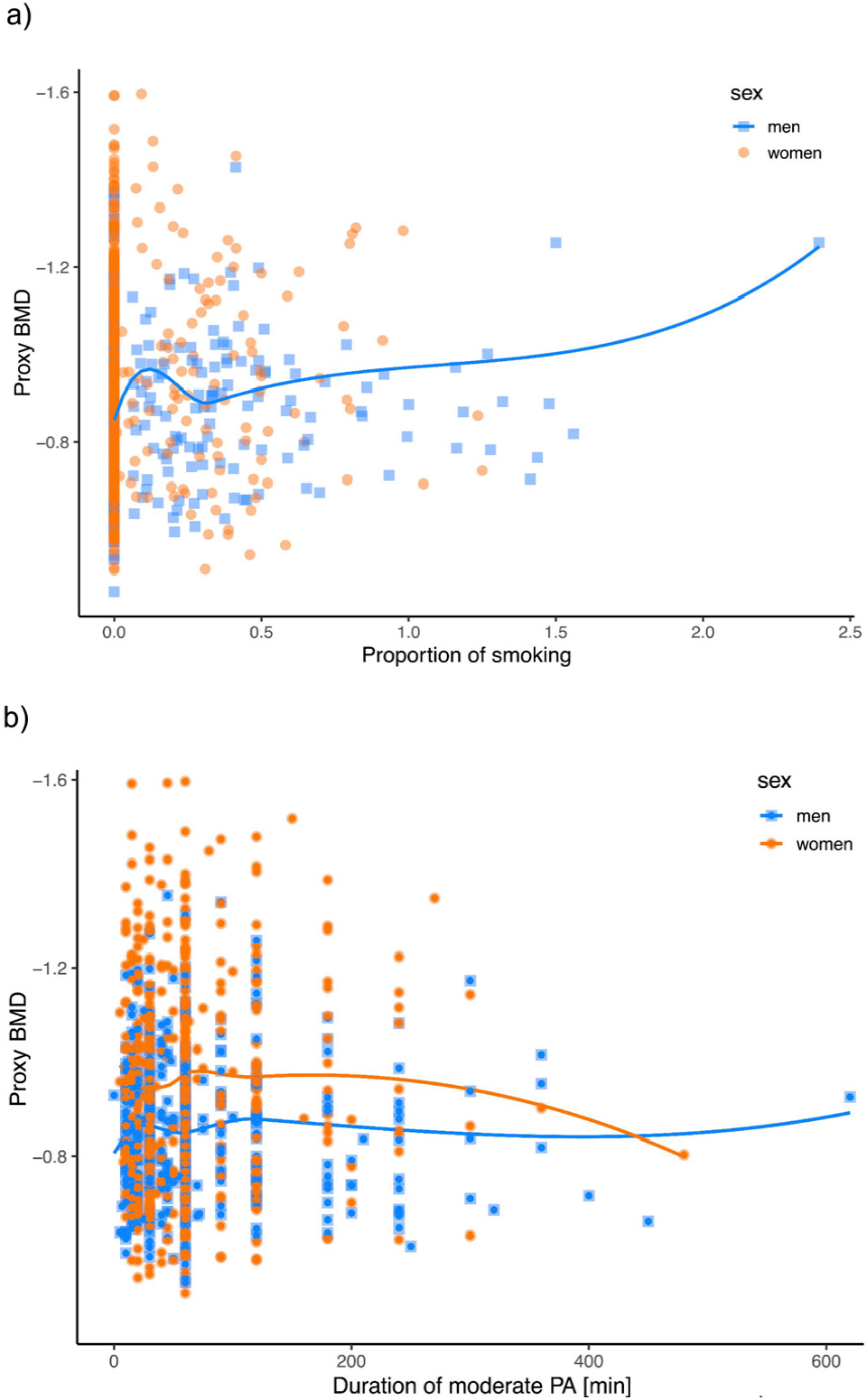
The association of proxy BMD measure and **a)** the amount of smoking and **b)** duration of physical activity for men and women. The negative effects of smoking are more evident in men, whereas positive effects of moderate physical activity are more prominent in women.

**Figure S5.**
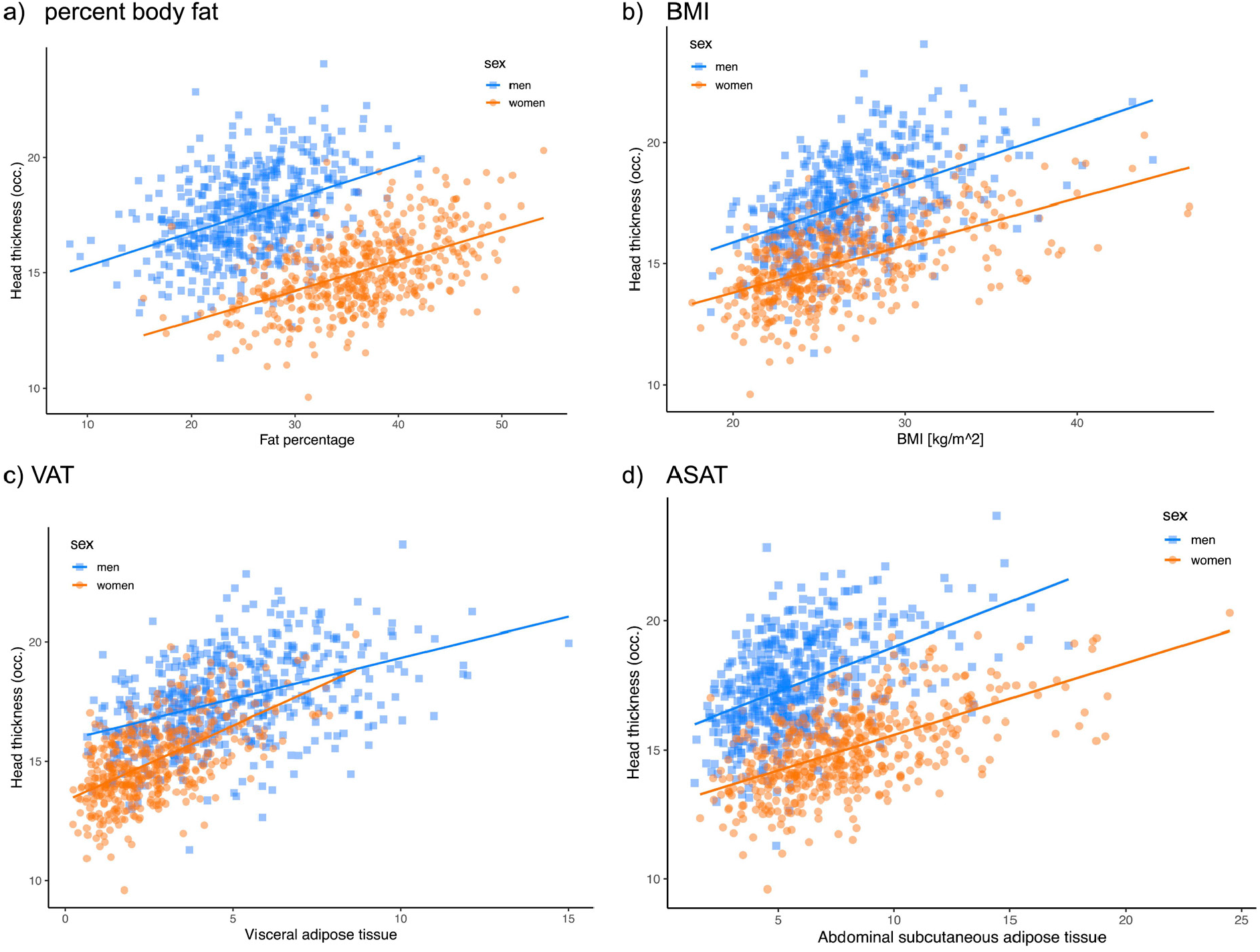
The association between head thickness estimation in the occipital part and **a)** percent body fat, **b)** BMI, **c)** VAT (visceral adipose tissue), and **d)** ASAT (abdominal subcutaneous adipose tissue).

**Figure S6.**
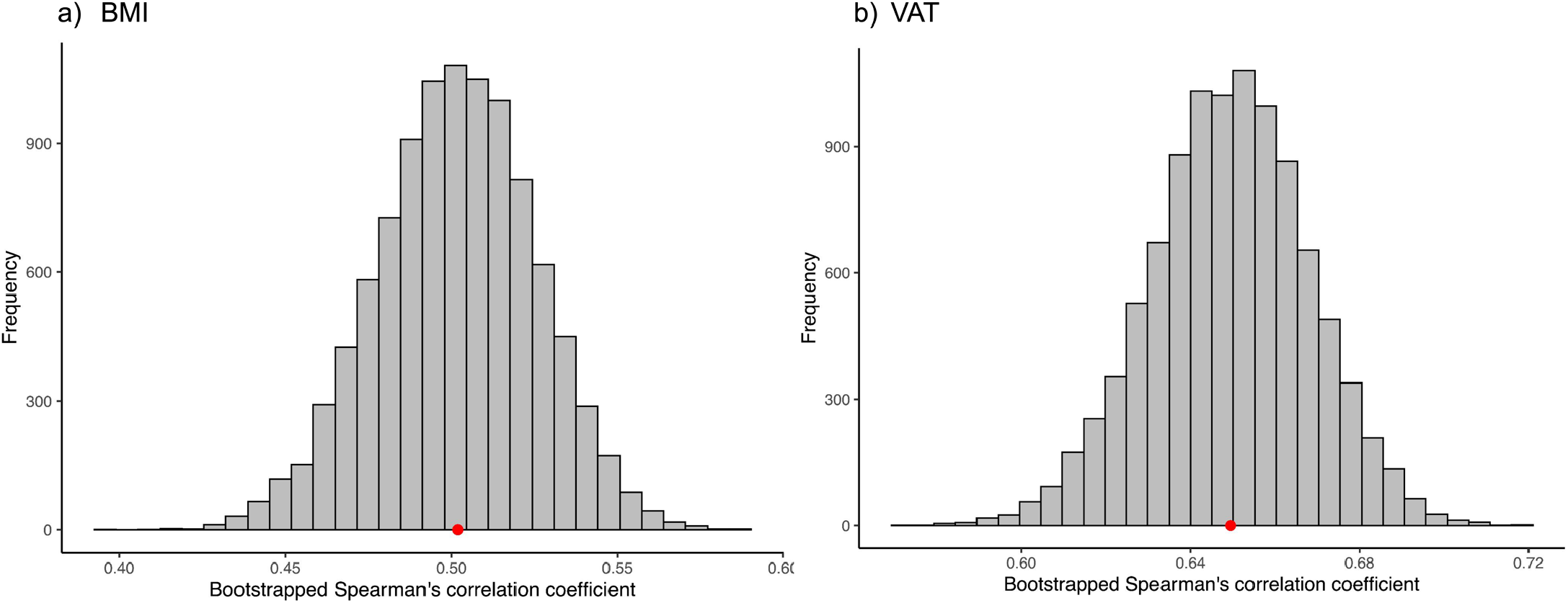
Bootstrapped Spearman’s correlation coefficients between the head thickness approximation and the a) BMI (rho = 0.50, p<.001, 95% CI: 0.45–0.55) and b) VAT (rho = 0.65, p<.001, 95% CI: 0.61–0.69).

